# Exploring *T. cruzi* IMPDH as a promising target through Chagas Box screening and AVN-944 inhibition

**DOI:** 10.1101/2025.08.06.668849

**Authors:** Lobo-Rojas Angel, Marchese Letícia, G. Eufrásio Amanda, D. T. Souza Gabriel, F. Catelli Michelle, D.N. Faria Jessica, T. Cordeiro Artur

## Abstract

Chagas disease, caused by *Trypanosoma cruzi*, remains a leading cause of heart failure in Latin America, with current treatments limited to acute-phase efficacy, significant toxicity, and emerging resistance. Inosine monophosphate dehydrogenase (IMPDH) is an essential enzyme in guanine nucleotide salvage pathway and represents a promising alternative target. Here, we combined computational screening, biochemical and cell-based phenotypic assays that support *T. cruzi* IMPDH (*Tc*IMPDH) as a druggable target and identify repurposing opportunities among clinical-stage inhibitors. Using Tanimoto similarity scoring against the library of 222 Chagas Box compounds, we identified TCMDC-143376 as uniquely similar to the clinical IMPDH inhibitors Merimepodib and AVN-944. Phylogenetic analysis and multiple sequence alignment confirmed conservation of both catalytic and allosteric residues—drawn from *T. foetus* and *T. brucei* structures—within *Tc*IMPDH. Recombinant *Tc*IMPDH kinetics revealed Michaelis constants of 155 µM for IMP and 292 µM for NAD⁺. Biochemical IC₅ ₀ assays showed submicromolar inhibition by AVN-944 (0.20 µM), (S)-Merimepodib (0.21 µM), and (R)-Merimepodib (0.37 µM). In H9c2 cardiomyoblasts infected with intracellular amastigotes, AVN-944 achieved the lowest EC₅ ₀ (0.4 µM) and highest selectivity index (SI > 100), outperforming benznidazole (EC₅ ₀ = 3.0 µM; SI = 13.4) and other inhibitors. Our findings support *Tc*IMPDH as a promising alternative drug target for Chagas disease and position AVN-944 as a compelling candidate to evaluate this therapeutic strategy in animal models.

## 1. Introduction

Chagas disease, caused by the protozoan parasite *Trypanosoma cruzi*, remains a major neglected tropical disease more than a century after its discovery [1]. Endemic in Latin America and increasingly detected in non-endemic regions due to migration [2], Chagas disease affects an estimated 6–7 million people worldwide [3]. Despite its public health burden, therapeutic options remain limited to just two frontline drugs: benznidazole and nifurtimox, both introduced over 50 years ago [4].

Benznidazole, the first-line treatment, induces DNA damage in *T. cruzi*; nonetheless, it has been observed the persistence of rare, non-replicative amastigotes in infected tissues [5]. Indeed, resistance to benznidazole can develop readily within a single population, by independently acquiring mutations in the gene encoding a mitochondrial nitroreductase which can give rise to distinct drug-resistant clones within a single population [6]. Nifurtimox, a nitrofuran prodrug, has been used for over 40 years to treat Chagas disease. Its mechanism of action was initially linked to oxidative stress, but newer evidence points to activation by a parasite-specific type I nitroreductase [7]. The resulting metabolite, a reactive open-chain nitrile, lacks selectivity, affecting both parasite and host cells, producing important side effects. Despite their different mechanisms, both nifurtimox and benznidazole are mainly effective in the acute phase. However, due to limited diagnostics and healthcare infrastructure in endemic regions, infections often go unnoticed until the chronic phase, where current drugs show poor efficacy. This underscores the urgent need for new therapeutic targets and treatments.

Alternative drug targets have been explored, including *T. cruzi* CYP51, a sterol 14α-demethylase essential for parasite viability [8,9]. Despite initial promise, clinical trials targeting CYP51 were halted due to suboptimal efficacy [10] and potential host toxicity [11]. Cytochrome b, a component of complex III in the mitochondrial electron transport chain, also has emerged as a druggable target in *Trypanosoma cruzi* and *Leishmania spp.* by phenotypic screening assays, with inhibitors like GNF7686 acting at the Qi site, blocking respiration and parasite growth [12]. However, its mitochondrial maxicircle-encoded nature makes it highly prone to resistance through point mutations, such as the L198F mutation in GNF7686-resistant *T. cruzi* [13]. Due to this high resistance potential, cytochrome b is now deprioritized in some drug discovery pipelines using resistant strain counterscreens [14,15]. On the other hand, proteasome inhibitors have emerged as promising anti-kinetoplastid agents, with compounds like GNF6702 demonstrating unprecedented *in vivo* efficacy, successfully clearing parasites in multiple mouse models [16]. This azabenzoxazole derivative acts as a non-competitive inhibitor of the kinetoplastid proteasome and laid the groundwork for potent analogs such as LXE408, a next-generation inhibitor characterized by high-resolution cryo-EM in complex with *Leishmania tarentolae* proteasomes, has progressed to clinical trials [17]. Despite these advances, challenges related to toxicity and selectivity remain, and no proteasome inhibitor has yet reached widespread clinical use for Chagas disease. These limitations highlight the continued need to explore alternative drug targets with safer and more effective therapeutic profiles.

One promising avenue lies in the purine salvage pathway, which is essential for *T. cruzi* survival due to its inability to synthesize purines *de novo* [18,19]. Instead, Trypanosomatids rely on salvaging purines from external sources, for instance, it has been previously observed that epimastigotes uptake hypoxanthine from LIT (Liver Infusion Tryptose) medium during its exponential growth phase [20]. In this pathway participate enzymes such as adenine phosphoribosyltransferase (APRT), hypoxanthine-guanine phosphoribosyltransferase (HGPRT), inosine-5′-monophosphate dehydrogenase (IMPDH) and Guanosine monophosphate synthase (GMPS), the first three enzymes have been identified in proteomic studies in *T. cruzi* glycosomes [21] and the last one has been demonstrated to be essential in the related organism *T. brucei* [22]. In particular, IMPDH plays a critical role by catalyzing the NAD⁺ -dependent oxidation of inosine monophosphate (IMP) to xanthosine monophosphate (XMP), the key precursor for guanine nucleotide biosynthesis [23]. While mammals possess IMPDH enzymes as well, parasite-specific differences in structure and regulation may offer opportunities for selective inhibition. It is worth noting that IMPDH has been explored as drug target in pathogenic prokaryotic organisms, it has been validated as a promising drug target in pathogens like *Helicobacter pylori* [24,25], *Bacillus anthracis*, *Staphylococcus aureus* and *Listeria monocytogenes* [26]. Structural determinants of inhibitor selectivity have been identified in bacterial IMPDHs [24]. Even in another protozoan parasite, the apicomplexan *Cryptosporidium parvum*, IMPDH has also been studied as a drug target [27]. The *C. parvum* IMPDH is closely related to the prokaryotic counterparts [28] and efforts are underway to repurpose *Cryptosporidium* IMPDH inhibitors as broad-spectrum antimicrobials [26].

IMPDH inhibitors for clinical use have been developed and explored for therapeutic purposes, ranging from immunosuppressants to antivirals and anticancer agents. Mycophenolic acid (MPA), an FDA-approved non-competitive IMPDH inhibitor is widely used as an immunosuppressant to prevent organ transplant rejection [29]. By selectively inhibiting IMPDH, interferes with the *de novo* synthesis of guanosine nucleotides, while most cell types can compensate via purine salvage pathways, T and B lymphocytes are particularly dependent on *de novo* purine synthesis, making them susceptible to IMPDH inhibition, as a result, MPA exerts potent cytostatic effects on lymphocytes, suppresses antibody production, and interferes with glycoprotein glycosylation involved in immune cell adhesion [30,31]. Its FDA-approved prodrug, Mycophenolate mofetil, offers improved oral bioavailability, avoiding some side effects but retaining the same mechanism of action [32]. Noteworthy, clinical evidence shows that Mycophenolate mofetil increases the risk of Chagas disease reactivation in heart transplant patients [33], also corroborated in mice revealed that although it reduces parasitemia, it does not improve survival during acute *T. cruzi* infection [34]. Another FDA-approved drug, ribavirin, exerts antiviral effects partly through IMPDH inhibition, but also directly targeting viral RNA polymerases [35]. More recent efforts have yielded second-generation IMPDH inhibitors such as VX-148, a potent IMPDH inhibitor [36] whose derivatives, AVN-944 and Merimepodib have progressed to clinical trials phase II. AVN-944, originally developed for cancer due to its capacity to deplete guanine nucleotides, inhibits ribosomal RNA synthesis and leads to nucleolar disruption, inducing apoptosis via cellular stress responses [37,38]. Similarly, Merimepodib, initially developed as an immunosuppressant [39], has also demonstrated broad-spectrum antiviral activity, including efficacy against Zika virus [40] and hepatitis C virus [41], further underscoring the therapeutic versatility of IMPDH inhibitors. Lastly, Mizoribine, a ribavirin derivative, is under investigation for rheumatoid arthritis [42]. Collectively, these compounds highlight the clinical relevance of IMPDH inhibition across diverse therapeutic areas and reinforce the rationale for repurposing such agents for neglected diseases like Chagas disease.

In this study, we focus on the *T. cruzi* IMPDH as a druggable target, using a ligand-based drug discovery approach, we identified potential inhibitors within the Chagas Box compound collection, a curated, open-source library of chemically diverse, drug-like molecules with confirmed activity against *T. cruzi*, made publicly available to accelerate Chagas disease drug discovery [43]. Notably, we identified TCMDC-143376, which showed structural similarity to AVN-944 and Merimepodib, suggesting potential cross-species inhibitory effects. Biochemical and phenotypic assays confirmed that both AVN-944 and Merimepodib are potent inhibitors of *Tc*IMPDH and display significant antiparasitic activity against *T. cruzi*. These findings not only reinforce *Tc*IMPDH as a promising and previously unexplored drug target for Chagas disease but also position AVN-944 and Merimepodib as valuable proof-of-concept compounds for validating *Tc*IMPDH inhibition as a viable therapeutic strategy.

## 2. Materials and Methods

### 2.1. In silico similarity analysis between IMPDH inhibitors and Chagas Box compounds

To assess the potential structural similarity between known IMPDH inhibitors and compounds with activity against *Trypanosoma cruzi*, we performed a Tanimoto similarity analysis [44]. The objective was to identify Chagas Box hits that share a high degree of molecular similarity with inhibitors previously developed for bacterial and human IMPDHs: The compounds used as queries were prokaryotic IMPDH inhibitors as C64, C91, A110 [24], Mycophenolate mofetil [45] and human IMPDH inhibitors as Mizoribine, Merimepodib and AVN-944.

These molecules were used as queries to calculate Tanimoto index against the 222 compounds Chagas Box set [43], all of them with confirmed activity against *T. cruzi* intracellular amastigotes. Molecular structures were processed as SMILES strings and similarity calculations were performed using the RDKit cheminformatics library in Python [46]. Molecules were converted to Morgan fingerprints (radius = 2), and pairwise Tanimoto similarity indices were computed between each reference inhibitor and all Chagas Box compounds. Iterative Grubbs’ test (α=0.0001) was used to identify outliers inside each population [47] and plotted using GraphPad Prism

9.5.0.

### 2.2. Bioinformatics and phylogenetic analysis

The *IMPDH* gene from *T. cruzi* was identified in the kinetoplastid genomic resource database (https://tritrypdb.org/tritrypdb/app) under the code TcCLB.507211.40, the translated sequence was retrieved as fasta format and used to perform a search by BLASTP (https://blast.ncbi.nlm.nih.gov/Blast.cgi) [48], restricted search sets were used by inclusion and/or exclusion of certain organisms or phylogenetic groups sequences to assure adequate sampling of *IMPDH* genes from landmark organisms. To perform the Multiple Sequence Alignment (MSA) IMPDH protein sequences from several organisms were used: *Trypanosoma cruzi* (XP_805772.1, which is the product from TcCLB.507211.40), *Trypanosoma brucei* (AAB46420.1) whose 3D structure was solved by following an *in cellulo* crystallization approach [49], *Leishmania donovani* (XP_003860332.1), two isoforms sequences from *Homo sapiens* 1 (NP_001136045.1) and 2 (NP_000875.2), two isoforms sequences from *Mus musculus* 1 (NP_001289862.1) and 2 (NP_035960.2), *Plasmodium falciparum* (XP_001352079.1), *Arabidopsis thaliana* (NP_178065.1), *Escherichia coli* (P0ADG7.1) and *Tritrichomonas foetus* (XP_068358496.1) whose x-ray solved structure is available at https://www.rcsb.org under the code 1MEW. Retrieved sequences in fasta format were aligned using the online program Clustal Omega [50], from the EMBL’s European Bioinformatics Institute (https://www.ebi.ac.uk/jdispatcher/msa/clustalo), and subsequently analyzed with the online server ESPript 3 (https://espript.ibcp.fr/ESPript/cgi-bin/ESPript.cgi) [51].

### 2.3. Cloning, heterologous expression and purification of the Trypanosoma cruzi IMPDH-product

The *IMPDH* gene from *Trypanosoma cruzi* was identified in the kinetoplastid genomic resource database (https://tritrypdb.org/tritrypdb/app) under the code TcCLB.507211.40, retrieved ORF sequence was sent to Genscript services in order to synthesize and clone into pET28a vector. Flanked with *Xho*I and *BamH*I restrictions sites. The sequence of the gene was optimized by Genscript, taking into account a wide variety of factors that regulate and influence gene expression levels, and by GenSmart codon optimization tool. Taking advantage of the restriction enzyme sites, the gene was subçloned into pETSUMO vector as previously made with other recombinant enzymes [52,53]. The resulting plasmid pETSUMO-*TcIMPDH* was used to transform BL21 by conventional procedures and overpressed by autoinduction [54] at 20 °C, 200 RPM for 72 hours. Bacterial cells were resuspended in buffer A (50 mM Tris-HCl pH 8.0 + 0.3 M KCl + 5 % glycerol) and clear lysate was obtained by sonication and centrifugation. Thus, it was charged onto a Ni-sepharose affinity column previously equilibrated with buffer A. After washing steps, the recombinant protein was eluted with increasing concentrations of buffer B (50 mM Tris-HCl pH 8.0 + 0.3 M KCl + 5 % glycerol + 1 M imidazole). Fractions containing the recombinant protein were pooled. His-SUMO tag was cleaved by incubation with ULP-1 protease at 4 °C overnight. Afterwards, a reverse affinity chromatography was made, and finally fractions with *Tc*IMPDH were pooled, added 10 % glycerol and 1 mM DTT, stored at -80 °C. Along purification steps, the protein concentration was assessed by Bradford method and the performance checked by SDS-PAGE [55].

### 2.4. Enzyme kinetic characterization

The Michaelis-Menten kinetic parameters (Km) for IMP and NAD⁺ were determined using the direct assay by following the formation of NADH fluorescence intensity (Ex = 340 nm; Em = 485 nm) at 25 °C, in black 384-wells microplates. Fluorescence was recorded every 30 seconds over 10 minutes using a CLARIOstar plate reader (BMG LabTech) and steady-state velocities were calculated from the maximum slope of the fluorescence curve. All reactions were performed in triplicate, under optimized buffer conditions: 50 mM tri-sodium citrate (pH 7), 100 mM KCl, and 1 mM DTT. For both substrates, 1 μM of purified recombinant *Tc*IMPDH was used. To determine the Km for IMP, NAD⁺ was held at a saturating concentration (1 mM), while IMP was tested in eight 1:2 serial dilutions starting from 1 mM. Conversely, to determine the Km for NAD⁺, IMP was fixed at 1 mM, and NAD⁺ was serially diluted in the same manner. Reactions were initiated by the addition of IMP, and data were analyzed using nonlinear regression to fit the Michaelis-Menten equation in GraphPad Prism 9.5.0.

### 2.5. Screening of Chagas Box library for TcIMPDH inhibitors

Compounds were assayed at a single concentration of 10 μM. They were provided by GSK (GlaxoSmithKline) in a 384-well ready-to use assay microplate containing 25 nL of each sample at 10 mM (100% DMSO). Because some tested compounds showed fluorescence at wavelengths E_x_ 340 nm /E_m_ 460 nm which are used to detect direct product NADH of the reaction, it was used the coupled assay with diaphorase and resorufin fluorescence (E_x_ 545 nm /E_m_ 600 nm) [52] and recorded every 30 s for 10 min. Velocities at steady state were calculated in the plate reader (CLARIOstar, BMG LabTech) as the maximum slope. Next, 20 μL of Mix solution containing 1.25 μM *Tc*IMPDH, 0.250 mM NAD^+^, 10 μM resazurin in 62.5 mM citrate buffer pH 7.0, 125 mM KCl and 1 mM DTT, 1.25 U/mL diaphorase was dispensed into wells and the reaction initiated by addition of 5 μL of 0.25 mM IMP. After 15 min, resorufin fluorescence (Ex = 570 nm; Em = 590 nm) was measured in a plate reader (CLARIOstar, BMG LabTech). Data were normalized and plotted using GraphPad Prism 9.5.0.

### 2.6. Measuring of half-maximal inhibitory concentration of compounds against TcIMPDH activity

To determine the concentration necessary to inhibit 50% of *Tc*IMPDH activity (IC_50_), stock solutions (10 mM) of compounds Mycophenolate mofetil, (S)-Merimepodib, AVN-944 and (R)-Merimepodib were serially diluted in DMSO, applying a 1⁄2-log dilution factor. The above mentioned diaphorase coupled assay was used. The final concentrations of components in the enzymatic assay were: 0.2 mM NAD^+^, 0.2 mM Inosine 5′-monophosphate (IMP), 1 μM *Tc*IMPDH, 1 U/mL diaphorase, 10 μM resazurin in 50 mM citrate buffer pH 7.0 plus 100 mM KCl. The reaction was started by adding IMP. *Tc*IMPDH activity was normalized, and the IC_50_ was calculated using GraphPad Prism 9.5.0.

### 2.7. Determining of the half-maximal effective concentration (EC₅₀) against Trypanosoma cruzi amastigotes in H9c2 cells

To determine EC_50_ to kill intracellular amastigotes in H9c2 host cells, we used an image-based assay previously described in [20]. Briefly, *T. cruzi* trypomastigotes (from LLC-MK2 supernatant) were used to infect H9c2 rat cardiomyocytes overnight. After washing, cells were incubated for 48 h in fresh DMEM, then seeded (1.5 × 10^3^ cells / well) into 384-well plates with 50 µL assay volume (0.4% DMSO) containing 0.2 µL of compounds. After 72 h, cells were washed, fixed with 4% paraformaldehyde, and stained with Hoechst 33342 (4 µg/mL). Images (5 per well) were acquired using an Operetta microscope (20X objective) and analyzed with Columbus software. Data on cell count and infection ratio were processed in GraphPad Prism 9.5.0.

## 3. Results

### 3.1 In silico identification of putative IMPDH inhibitors in the Chagas Box collection

To find structural similarities between IMPDH inhibitors and *Trypanosoma cruzi* active compounds in the Chagas Box, we compared the SMILES of prokaryotic IMPDH inhibitors (A110, C64, C91), or human IMPDH inhibitors (Mycophenolate, Mizoribine, Merimepodib and AVN-944) –showed in Fig. 1A– with the Chagas Box library using Tanimoto similarity scoring (Fig. 1B). IMPDH prokaryotic inhibitors yielded no significant Tanimoto scoring with any of the Chagas Box compound set, while when using Merimepodib and AVN-944, the analysis revealed that only **TCMDC-143376** exhibited Tanimoto indexes of 0.817 and 0.506 for any enantiomer of Merimepodib and AVN-944, respectively. Such scores were statistically different (α = 0.0001) of those displayed by the rest of the Chagas Box molecules. This result highlights the unique structural resemblance of TCMDC-143376.

**Fig. 1.**
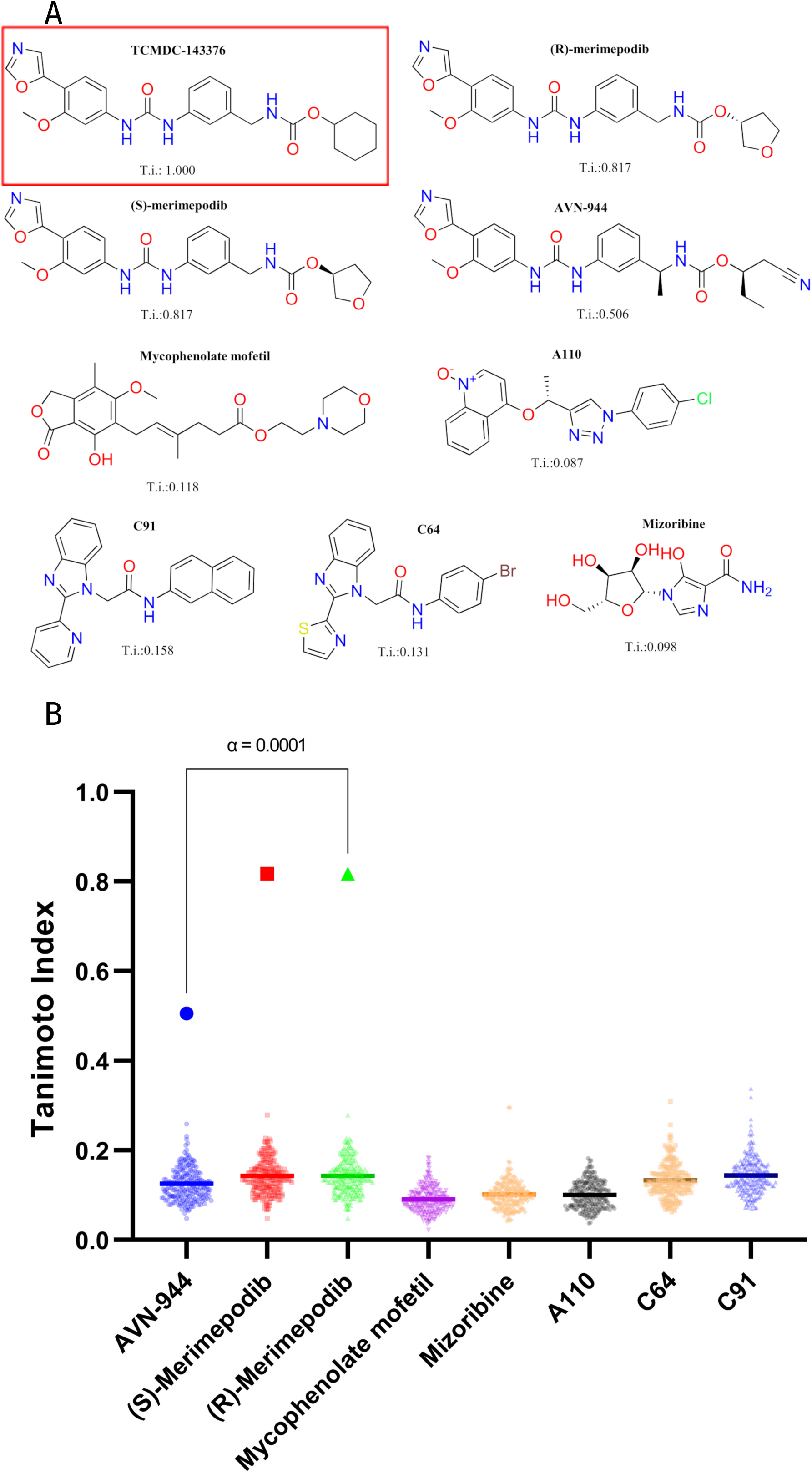
Structures of IMPDH inhibitors and structural similarity analysis of Chagas Box compounds. **(A)** Chemical structures of the reference inhibitors used in the analysis, including their corresponding Tanimoto similarity scores relative to the compound TCMDC-143376 (phenotypic hit from Chagas Box, enclosed with a red rectangle). **(B)** Tanimoto similarity indices of 222 Chagas Box compounds, computed against a set of known IMPDH inhibitors: AVN-944, (S)-Merimepodib, (R)-Merimepodib, Mycophenolate mofetil, Mizoribine, and the prokaryotic inhibitors A110, C91, and C64. Iterative Grubbs’ method with an α=0.0001 to identify outliers inside each sample was used (GraphPad Prism 9.5.0).

A phylogenetic analysis performed using IMPDH sequences from across the tree of life (**see Fig. S1**) revealed that IMPDHs from Trypanosomatidae cluster with those from Chordata and Fungi. This evolutionary proximity corroborates the observed *in silico* cross-species structure similarities of inhibitors such as AVN-944 and Merimepodib. In contrast, most Apicomplexan IMPDHs form a distinct clade within eukaryotes, separate from their prokaryotic counterparts, with the notable exception of the *Cryptosporidium* genus, whose IMPDH sequences group within the bacterial cluster, as previously reported [28,56].

It is likely that TCMDC-143376 may be targeting *T. cruzi* IMPDH. This structural insight provided a strong rationale for further investigating *Tc*IMPDH as the potential molecular target of TCMDC-143376, and by extension, for exploring the entire Chagas Box library in the context of IMPDH inhibition. Therefore, we proceeded to study, clone and recombinantly express *Trypanosoma cruzi* IMPDH to enable further biochemical and pharmacological characterization of this enzyme as a potential drug target.

### 3.2 Molecular and biochemical characterization of TcIMPDH

The *IMPDH* gene from *Trypanosoma cruzi* was identified in the kinetoplastid genomic resource database (https://tritrypdb.org/tritrypdb/app) under the code TcCLB.507211.40 located in chromosome 8. In the *T. cruzi* genome exists another copy of this gene, located in chromosome 40 with a 98 % identity, with accession code TcCLB.511351.9. The *Tc*IMPDH gene is a 1539-bp long open-reading frame which encodes for a polypeptide of 512 amino acids with a predicted molecular mass of 55.6 kDa. Both protein isoforms have been identified in proteomic studies of *Trypanosoma cruzi* glycosomes, consistent with the presence of a canonical peroxisomal targeting signal type 1 (PTS1, SKL) in both sequences [21]. IMPDH is a member of the Pfam00571 family, characterized by the IMPDH/GMP reductase domain, which adopts a canonical TIM barrel structure (α/β-barrel), composed of 8 α-helices and 8 parallel β-strands [57]. In addition to the catalytic core, two CBS (cystathionine beta-synthase) domains are inserted within the barrel, often associated with allosteric regulation as found elsewhere for other IMPDHs and potentially occurring in Trypanosomatid IMPDHs as it was observed *T. brucei* IMPDH CBS domain interacting with GMP and ATP *in cellulo* crystals [49].

Based on the structure of *T. foetus* IMPDH in complex with XMP and NAD^+^ [58], the binding sites of this structure were mapped in *Tc*IMPDH. XMP/IMP site is defined by a conserved set of residues: S317, D358, E408, G409, E431. Through alignment (see Fig. S2), we identified the equivalent residues in *T. cruzi* IMPDH as: **S323**, **D358**, **M408** (instead of E as in *T. foetus*), **G409**, **Q435** (replacing E found in *T. foetus*). In relation to the NAD⁺ binding residues identified in *T. foetus* [58] are: G314, D261, S262, S263, R241, W269. In *T. cruzi*, the equivalent residues are: **G320, D268, S269, S270, R247, Y276**. The conservation of core catalytic residues across bacteria, protozoan and mammalian species supports a shared catalytic mechanism, also observed by other authors [59].

In addition to the conservation of catalytic residues, alignment-based comparison between *T. cruzi* and *T. brucei* IMPDH—proteins sharing 81% sequence identity—provides insight into the regulatory Bateman domains. Structural studies of *T. brucei* IMPDH (*Tb*IMPDH) obtained from *in cellulo* crystals [49] identified a GMP binding site within this domain, involving residues **K115**, **S136**, and **G137**. These same residues are conserved in *T. cruzi* IMPDH (*Tc*IMPDH) at identical positions (see Fig. S2), suggesting functional conservation. Furthermore, the ATP-binding site in *Tb*IMPDH is composed of **S136**, **T156**, **K157**, **D158**, **T174**, **T180**, **H200**, **Y202**, and **R219**. In *Tc*IMPDH, all these residues are conserved, with the exception of a single conservative substitution of T156 by a serine residue. This high degree of conservation reinforces the structural and functional similarities between these ortholog enzymes.

To enable further analysis of *Tc*IMPDH, the purified recombinant enzyme was kinetically characterized. Steady-state activities were fitted to the Michaelis-Menten model to derive kinetic parameters. The resulting Km values were **155 μM** (95 % confidence interval 113.3–197.8 μM) and **292 μM** (95 % confidence interval 221.6–364.0 μM) for IMP and NAD⁺, respectively (Fig. S3). Compared to *T. brucei* IMPDH (Km(IMP) = 30 μM, Km(NAD⁺) = 1300 μM) [60] and *L. donovani* IMPDH (Km(IMP) = 33 μM, Km(NAD⁺) = 390 μM) [61], *Tc*IMPDH shows a moderately lower affinity for IMP but an intermediate affinity for NAD⁺. These values also contrast sharply with human IMPDHs, which exhibit much tighter binding to both substrates (Km(IMP) = 4–18 μM; Km(NAD⁺) = 6–70 μM) [62].

### 3.3 Screening of the Chagas Box

To validate the *in silico* predictions, we screened the Chagas Box collection for *Tc*IMPDH inhibitors., which was kindly provided by GSK in a format of ready-to-use 384-well black plates (for more details see materials and methods section). The screening assay employed the coupled reaction system, in which *Tc*IMPDH activity was linked to diaphorase-mediated resazurin reduction as previously described [52]. The primary screening, conducted in two independent runs (Fig. 2), consistently identified a single compound as a putative *Tc*IMPDH inhibitor: TCMDC-143376. This result corroborates our *in silico* prediction. This compound consistently showed 40–45% inhibition across two independent runs, suggesting an IC_50_ below 10 μM (the fixed concentration used for all compounds in the Chagas Box screening). To validate inhibitory activity, GSK kindly provided resupplied samples, enabling further confirmatory assays.

**Fig. 2:**
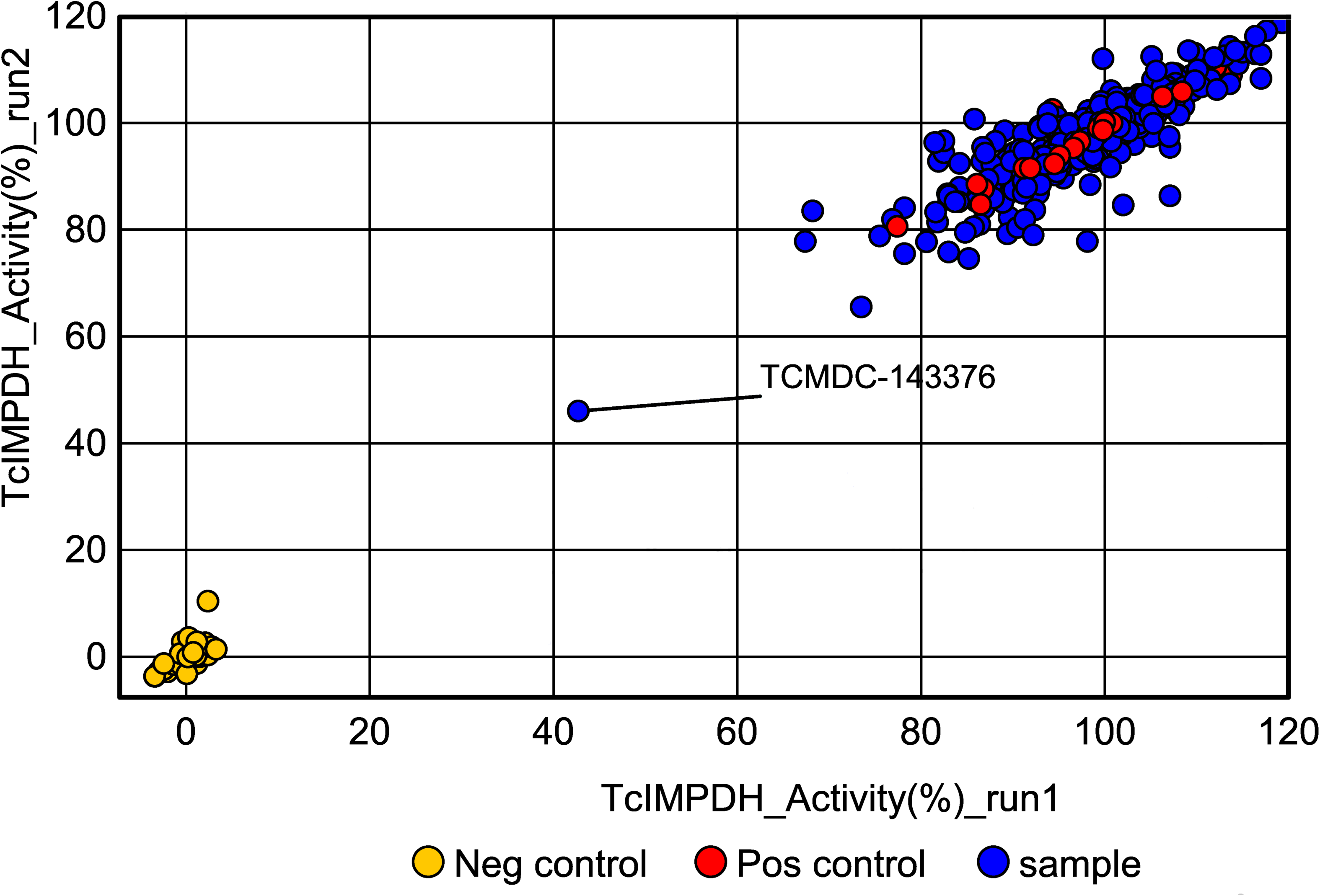
Chagas Box screening for identification of *Tc*IMPDH modulators. The experiment was performed in duplicate and *Tc*IMPDH activity in the presence of each sample was normalized using negative control reactions without addition of the IMP substrate (0% activity) and positive reactions without compounds (100% activity).

### 3.4 Confirming TCMDC-143376 as an IMPDH inhibitor and assessing other compounds against T. cruzi IMPDH

Half-maximal inhibitory concentrations (IC₅₀) were measured to confirm the inhibition of *Tc*IMPDH by TCMDC-143376 and assess other structural similar compounds, including Mycophenolate Mofetil, AVN-944 and Merimepodib enantiomers. They showed inhibitory activity against *Tc*IMPDH with IC₅ ₀ values in the low micromolar to submicromolar range. TCMDC-143376 exhibited an IC₅ ₀ of 4.00 µM, comparable to that of Mycophenolate Mofetil (4.40 µM), while AVN-944 and (S)-Merimepodib demonstrated much stronger inhibition, with IC₅ ₀ values of 0.20 µM and 0.21 µM, respectively. (R)-Merimepodib showed slightly reduced potency with an IC₅ ₀ of 0.37 µM. Consistent with *in silico* results, Mizoribine did not show measurable inhibition of *Tc*IMPDH activity under the conditions tested. These results confirm the susceptibility of *Tc*IMPDH to structurally similar IMPDH inhibitors and support the utility of AVN-944 and Merimepodib enantiomers as potential hit compounds.

### 3.5 In vitro activity of IMPDH inhibitors against Trypanosoma cruzi amastigotes into H9c2 host cells

To evaluate the *in vitro* therapeutic potential of known IMPDH inhibitors against *Trypanosoma cruzi* amastigotes, we assessed their Half-maximal efficacy concentration (EC₅₀), cytotoxicity (CC₅₀), and the consequent selectivity index (SI = CC₅ ₀ /EC₅₀) was also calculated in an image-based assay in H9c2 rat cardiomyoblasts as previously described [20]. Representative images are shown in Fig. 3 with the respective close-ups underscoring details of image-based assays, by comparing the same concentration (1.2 µM) it is noted that AVN-944 bears higher potency than the first-line treatment drug: Benznidazole. Among the tested compounds, AVN-944 exhibited the highest potency (EC₅ ₀ = 0.4 µM) and the most favorable selectivity index (SI = >100), followed by benznidazole (SI = 13.4) and Mycophenolate Mofetil (SI = 6.6). While (S)-Merimepodib and its (R)-enantiomer also showed activity, they exhibited lower selectivity, largely due to their moderate toxicity in H9c2 cells (see table 2). Similar selectivity indexes and cytotoxicity have been obtained for AVN-944 when studied against Foot-and-mouth disease virus or Mpox virus [63,64] and for Merimepodib other authors also found cytotoxic effects even when host cells are guanosine-rescued [40,63].

**Fig. 3.**
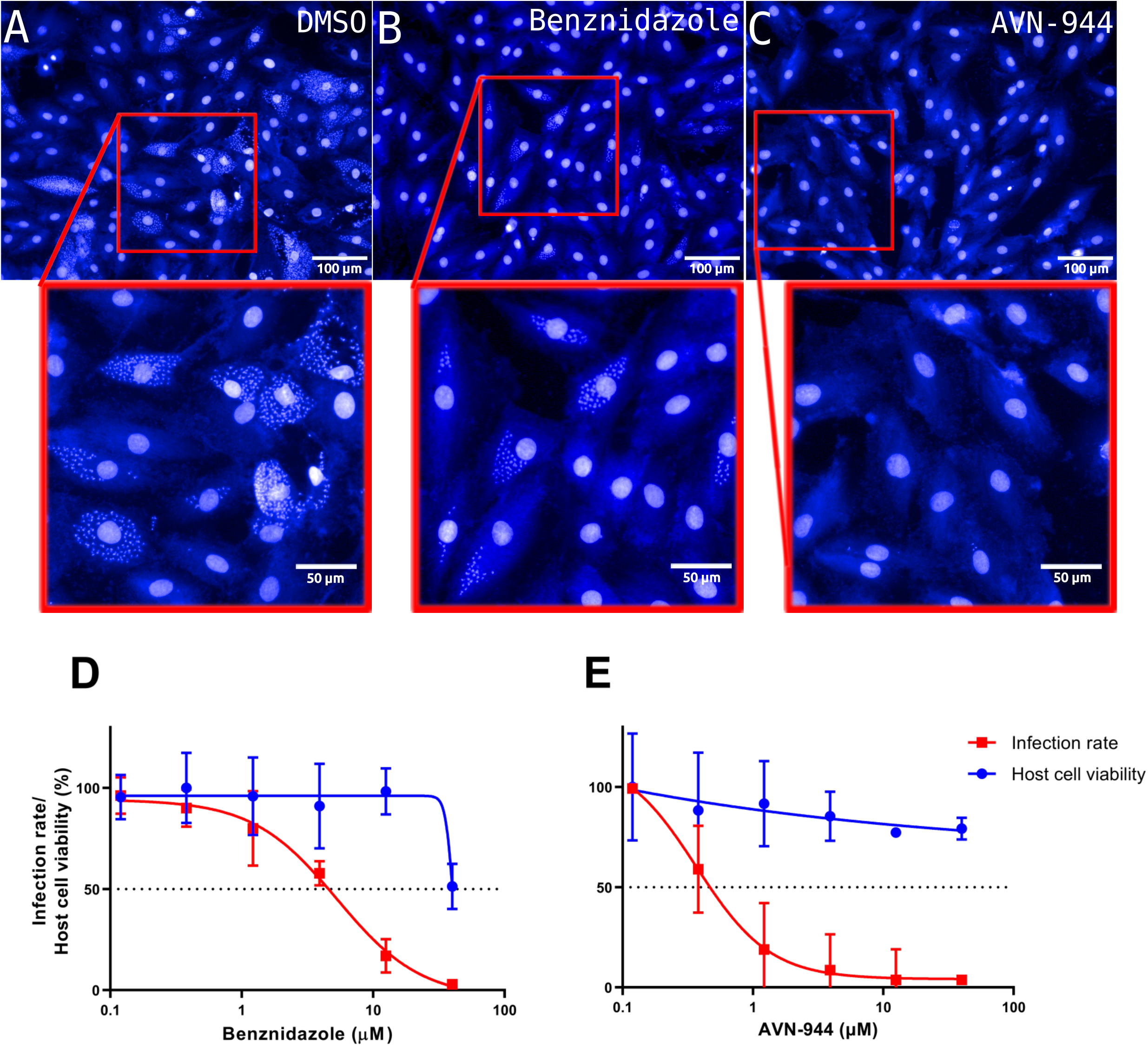
*In vitro* effect of benznidazole and AVN-944 on *Trypanosoma cruzi* amastigotes in H9c2 cells. H9c2 cardiomyoblast cells were infected with *T. cruzi* trypomastigotes, washed to remove extracellular parasites, and then incubated with the corresponding compound. Representative micrographs are shown for each condition (A) vehicle control (DMSO), (B) benznidazole (1.2 μM), or (C) AVN-944 (1.2 μM), accompanied by close-up images highlighting intracellular amastigotes. Panels (D) and (E) depict dose-response curves fitted to a four-parameter logistic model for benznidazole and AVN-944, respectively. The normalized infection rate (red) and host cell viability (blue) are plotted with corresponding standard deviations (N ≥ 3).

**Table 1.**
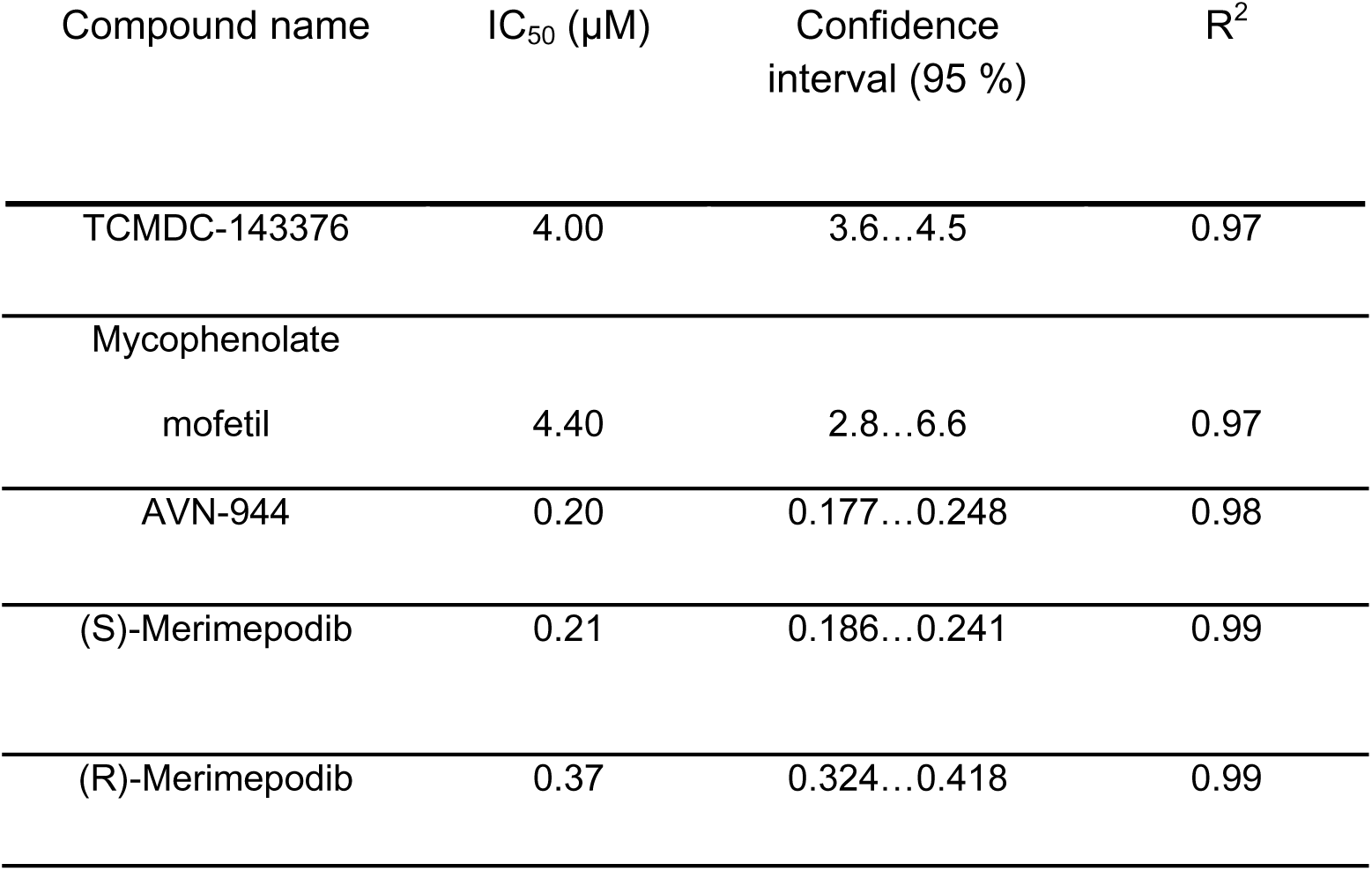
IC₅ ₀ values of TCMDC-143376 (from Chagas Box) and other known IMPDH inhibitors against *Trypanosoma cruzi* IMPDH. Compounds were tested in triplicate with 10-point serial dilutions in the presence of 0.2 mM IMP and 0.2 mM NAD⁺. IC₅ ₀ values and 95% confidence intervals were calculated using nonlinear regression in GraphPad Prism 9.5.0.

**Table 2.** EC₅ ₀, CC₅ ₀, and Selectivity Index (SI) of IMPDH inhibitors and benznidazole against *T. cruzi* amastigotes in H9c2 cells. Anti-amastigote activity (EC₅₀), host cell cytotoxicity (CC₅₀), and SI were determined from image-based infection assays. Data represent averages and standard deviations (SD) of at least two biological replicates.

| Compound name | EC<sub>50</sub> |  | CC<sub>50</sub> |  | SI |
| --- | --- | --- | --- | --- | --- |
|  | Average | SD | Average | SD |  |
| Benznidazole | 3.0 | 1.3 | ~40.2 | 6.4 | 13.4 |
| TCMDC-143376** | ~15.9 |  | ~100 |  | 6.29 |
| Mycophenolate mofetil | 1.5 | 1.0 | 9.9 | 6.9 | 6.6 |
| (S)-Merimepodib | 4.8 | 3.5 | 13.6 | 10.1 | 2.9 |
| (R)-Merimepodib | 3.0 | 1.5 | 9.1 | 7.7 | 3.0 |
| AVN-944* | 0.4 | 0.2 | >40 | 20.7 | >100 |

\*SI obtained for this compound is approximated due to uncertainty in CC<sub>50</sub> values.
\*\*Data extracted from [43] and obtained in Hepatoblastoma cell line (HepG2) as host cells.

For comparative purposes, we considered the previously reported activity of TCMDC-143376—originally annotated as a putative CYP51 inhibitor— [43]. Although its anti-amastigote potency was modest (∼15.9 μM), it showed favorable toxicity profiles in mammalian cell lines in HepG2 CC_50_ ≃ 100 μM. By contrast, AVN-944 demonstrated superior potency (EC₅₀ = 0.4 µM) suggesting a better therapeutic window.

Importantly, host-cell cytotoxicity (CC₅₀) was determined by counting total H9c2 cells after 72 h of compound exposure, which also quantifies intracellular amastigotes. Over this period, untreated cardiomyoblasts proliferate, so control wells show increased cell numbers; it also occurred in benznidazole-treated cells, which do not inhibit host IMPDH, host cells continued to proliferate and yielded higher cell counts. In contrast, IMPDH inhibitors block host IMPDH activity (indeed at lower concentrations), arresting cell division without inducing death; cells remain viable but do not multiply [65-67]. To distinguish this cytostatic effect from true cytotoxicity, CC₅ ₀ values were normalized to the lowest tested compound concentration, leading to a better fit of the four-parameter logistic equation (see comparison in Fig. S4).

## 4. Discussion

In this study, we employed an integrated computational, biochemical, and cellular strategy to validate *Tc*IMPDH as an alternative therapeutic target for Chagas disease. Tanimoto-based similarity screening of known phenotypic hits from GSK Chagas Box identified TCMDC-143376 as structurally related to known human IMPDH inhibitors Merimepodib and AVN-944 [44]. Further *in silico* studies by phylogeny with IMPDH primary sequences revealed that Trypanosomatidae’s ones cluster with vertebrate and fungal orthologs, supporting previous results with Tanimoto indexing and giving insights about possible cross-species activity of these inhibitors. In light of these *in silico* results, we cloned, expressed, and kinetically characterized the *Tc*IMPDH, enabling biochemical validation. Indeed, TCMDC-143376 inhibited *Tc*IMPDH activity, consistent with our *in silico* approach, but showed moderate potency, comparable to Mycophenolate mofetil, while AVN-944, (S)-Merimepodib and (R)-Merimepodib were submicromolar inhibitors. Mizoribine was inactive, validating the computational prioritization. In infected H9c2 cardiomyoblasts, AVN-944 demonstrated the highest potency, outperforming the first-line treatment drug: benznidazole. Together, these results position IMPDH as a promising target for the development of drugs against Chagas disease, and by extension for other related diseases such as leishmaniasis and sleeping sickness.

Concretely IMPDH inhibition targets the bottleneck of the purine salvage pathway, which emerges as a promising route for therapeutic intervention. Despite the apparent redundancy of the guanine nucleotide salvage pathway in trypanosomatids (see Fig. 4)—where GMP can be synthesized either directly from guanine via HGPRT, or indirectly from hypoxanthine through the sequential actions of HGPRT, IMPDH, and GMPS—evidence suggests that the latter route is predominantly utilized. This is supported by the essentiality of GMPS in *Trypanosoma brucei*, as demonstrated by Li et al. (2015) [22], who found that GMPS-null parasites could not sustain growth unless provided with supraphysiological concentrations of guanine (100 µM), far exceeding those encountered in physiological environment. Moreover, hypoxanthine acted as a competitive inhibitor of guanine-mediated rescue, implying a shared uptake or metabolic channel between these purines. Since GMPS catalyzes the final conversion of XMP (the product of IMPDH) into GMP (see Fig. 4), these findings indicate that guanine uptake and conversion alone are insufficient under normal conditions to compensate for loss of the IMPDH-GMPS route (see Fig. 4). Thus, despite theoretical redundancy, the metabolic flux in trypanosomatids heavily favors GMP production via hypoxanthine salvage through IMPDH and GMPS, reinforcing the critical role of these enzymes in nucleotide homeostasis and parasite viability.

**Fig. 4.**
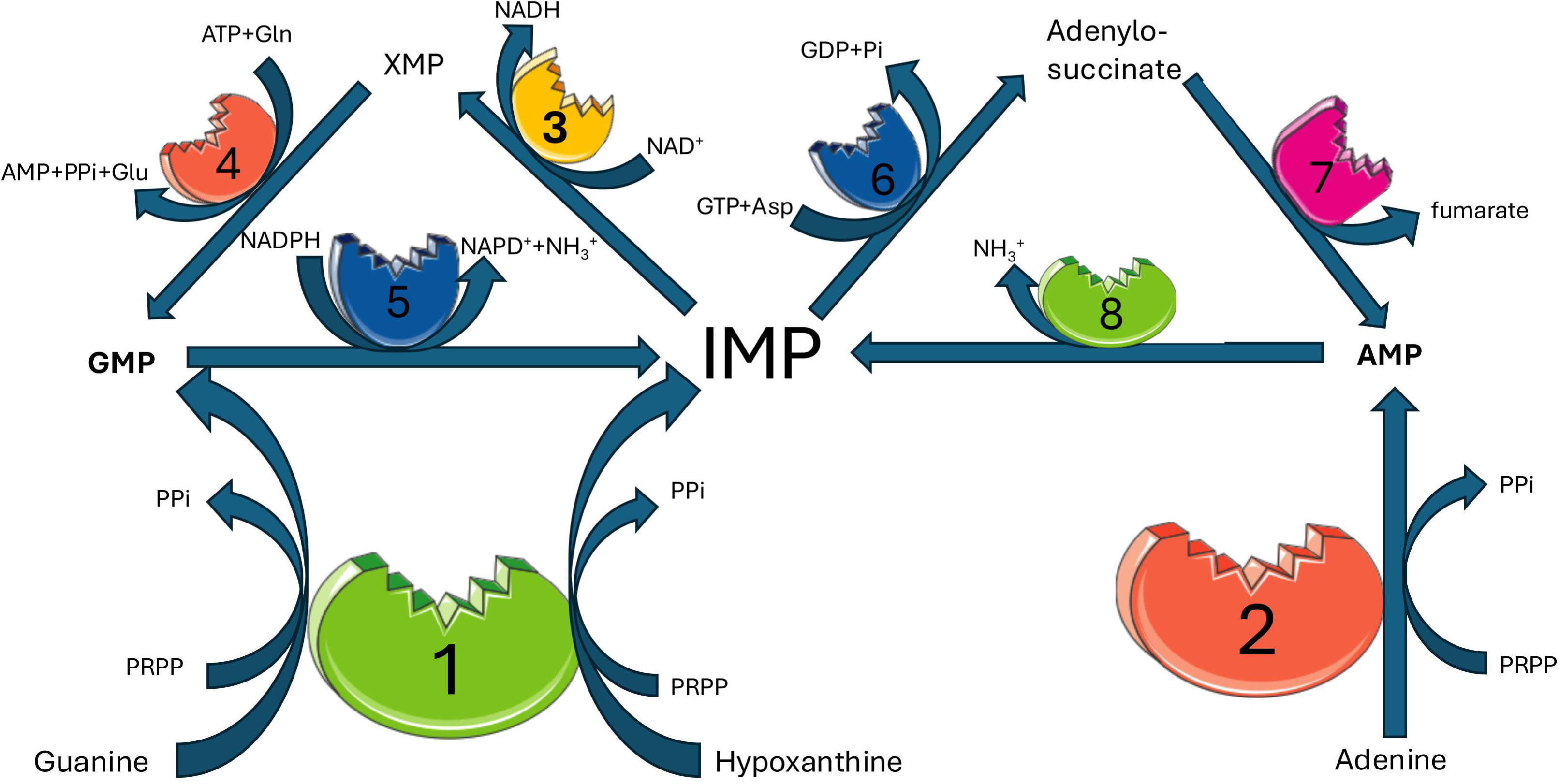
Schematic representation of purine salvage metabolism in *Trypanosoma cruzi* showing the context of IMPDH role. IMP plays a central role, being derived for the production of AMP or GMP, recycling steps catalyzed by 5 and 8 return GMP or AMP to IMP central pool. In Trypanosomatids, despite apparent redundancy for GMP production, it is mainly synthesized through the sequential action of 1, 3 and 4 from Hypoxanthine, rather than directly through 1 from Guanine. Enzymes: **1**: hypoxanthine-guanine phosphoribosyltransferase (HGPRT), **2**: adenine phosphoribosyltransferase (APRT), **3**: inosine-5-monophosphate dehydrogenase (IMPDH), **4**: GMP synthase (GMPS), **5:** GMP reductase (GMPR), **6**: adenylosuccinate synthase,**7**: adenylosuccinate lyase, **8**: AMP deaminase. **Abbreviations:** PRPP: phosphoribosyl pyrophosphate, PPi: pyrophosphate, IMP: Inosine-5-monophosphate, AMP: adenosine-5-monophosphate, GMP: guanosine-5-monophosphate, XMP: xanthosine-5-monophosphate, NAD^+^: nicotinamide adenine dinucleotide oxidized form, NADH: nicotinamide adenine dinucleotide reduced form, Gln: l-glutamine, Glu: l-glutamate, Asp: l-aspartate, Pi: orthophosphate.

AVN-944 exhibited potent antiparasitic activity in our *in vitro* model using H9c2 cardiomyoblasts infected with *T. cruzi*, significantly outperforming both benznidazole—the current frontline therapy—and Mycophenolate mofetil. Specifically, AVN-944 achieved the lowest EC₅ ₀ and the highest selectivity index among the compounds tested, indicating a strong therapeutic potential with a favorable safety margin in host cells. Beyond its *in vitro* potency, AVN-944 is already in phase II clinical trials for oncology indications, which suggests, despite not having been published, it may have undergone extensive pharmacokinetic and toxicological evaluation in humans. These encouraging results justify further investigation of AVN-944 in animal models as a proof of concept for targeting *T. cruzi* IMPDH, which may be extended for other trypanosomatids too. Nevertheless, other evidence must be taken into account: Mycophenolate mofetil has been tested in Chagas murine model showing the reduction of parasitemia, but it fails to improve survival outcomes or achieve parasitological cure [34]. Furthermore, clinical evidence has raised safety concerns, as Mycophenolate mofetil has been implicated in *T. cruzi* reactivation in immunosuppressed patients as part of the treatment to avoid rejection of heart transplantation [33].

The development of AVN-944 and Merimepodib were originally aimed at targeting human IMPDH isoenzymes, which explains their effective performance in cancer [65,68] and antiviral therapies [40,63,69]. However, these therapeutic benefits are often accompanied by immunosuppressive effects. In humans, IMPDH is a key enzyme in the *de novo* synthesis of guanine nucleotides, which are essential for DNA and RNA synthesis, particularly during immune cell activation and proliferation. Upon T cell activation, both IMPDH1 and IMPDH2 isoforms are upregulated, leading to a sharp increase in IMPDH activity that supports the rapid expansion required for an effective immune response [70-72]. This immunological context is highly relevant for Chagas disease, where a robust immune response is crucial. During the acute phase, both B and T lymphocytes undergo marked expansion [73,74]. The balance between effector and regulatory lymphocytes is also critical. For instance, regulatory T cells (Tregs) expand in individuals with the indeterminate (asymptomatic) form of Chagas disease, helping to limit tissue damage by controlling hyperactive immune responses [75]. Similarly, expansion of specific B cell subsets—such as CD11b⁺ B1 B cells—has been associated with improved cardiac function and a protective immune profile [76,77]. And finally In the chronic phase, CD4⁺ CD8⁺ double-positive T cells retain the ability to produce effector molecules against the parasite [78].

The limitations of current IMPDH inhibitors—particularly their immunosuppressive effects—highlight the need for more selective compounds. Although AVN-944 and merimepodib are known to act as non-competitive inhibitors of IMPDH [23,79], their exact binding site on the enzyme remains unidentified. Determining this site is crucial, as it would enable the rational design of new inhibitors that specifically target trypanosomatid IMPDHs while sparing the human isoforms. This selectivity is especially important not only to avoid side effects but also to preserve the host’s natural immune response, which plays a critical role in controlling *T. cruzi* infection and improving clinical outcomes [70-78]. While some structural insights have been gained from the recently solved *in-cellulo* crystal structure of *T. brucei* IMPDH [49], the binding pocket for these inhibitors has not yet been mapped. Identifying this region will be key to exploiting structural differences between human and parasite enzymes, and it will guide the repurposing of AVN-944 and Merimepodib scaffold for the development of safer, more effective chemotherapies that work in synergy with the host immune system.

## 5. Conclusions

This study identifies *Tc*IMPDH as a promising drug target for Chagas disease through an integrated *in silico*, biochemical, and *in vitro* cellular approach. Using ligand-based virtual screening, we linked the Chagas Box hit TCMDC-143376 to known human IMPDH inhibitors, notably AVN-944 and Merimepodib. Biochemical and kinetic validation confirmed *Tc*IMPDH inhibition, with AVN-944 displaying potent enzymatic and antiparasitic activity, outperforming benznidazole in infected cardiomyoblasts. Our results support the essentiality of guanine nucleotide salvaging via IMPDH in *T. cruzi*, despite the theoretical redundancy of the purine salvage pathway. Importantly, AVN-944’s submicromolar efficacy, combined with its favorable selectivity index and clinical-stage status, underscores its potential to perform the proof of concept of targeting IMPDH in animal models. However, its known immunosuppressive effects in humans highlight the need for more selective inhibitors that preserve host immune function. Mapping the binding site of AVN-944 on trypanosomatid IMPDH will be critical for the rational design of parasite-selective compounds with improved safety and efficacy profiles for Chagas disease and related kinetoplastid infections.

## Abbreviations

IMPDH: Inosine-5-phosphate dehydrogenase
*Tc*IMPDH: Inosine-5-phosphate dehydrogenase from *Trypanosoma cruzi*
SMILES: Simplified Molecular Input Line Entry System
GMP: Guanosine monophosphate
BLASTP: Basic Local Alignment Tool for Proteins

## Ethics approval and consent to participate

Not applicable.

## Authorship contribution statement

**Lobo-Rojas Ángel:** Conceptualization, Data curation, Formal analysis, Investigation, Methodology, Software, Validation, Visualization, Writing – original draft, Writing – review & editing. **Marchese Letícia:** Conceptualization, Data curation, Formal analysis, Investigation, Methodology, Validation, Visualization. **Eufrásio Amanda, G:** Investigation, Methodology. **Souza Gabriel D. T:** Investigation. **Catelli Michelle, F:** Investigation, Methodology, Project administration. **Faria Jessica D.N:** Investigation, Methodology. **Cordeiro Artur, T:** Conceptualization, Data curation, Formal analysis, Funding acquisition, Project administration, Resources, Supervision, Validation, Writing – review & editing.

## Acknowledgements

We thank Ramón Borges for his assistance in illustrating the molecular structures and to Celso Eduardo Benedetti for kindly revising the manuscript.

## Availability of data

All relevant data are included in the manuscript and Supplementary information provided.

## References

[1] K. C. F. Lidani et al., “Chagas Disease: From Discovery to a Worldwide Health Problem,” Front. Public Health, vol. 7, p. 166, Jul. 2019, doi: 10.3389/fpubh.2019.00166.

[2] E. E. Conners, J. M. Vinetz, J. R. Weeks, and K. C. Brouwer, “A global systematic review of Chagas disease prevalence among migrants,” Acta Trop., vol. 156, pp. 68–78, Apr. 2016, doi: 10.1016/j.actatropica.2016.01.002.

[3] Z. M. Cucunubá et al., “The epidemiology of Chagas disease in the Americas,” Lancet Reg. Health - Am., vol. 37, p. 100881, Sep. 2024, doi: 10.1016/j.lana.2024.100881.

[4] P. García-Huertas and N. Cardona-Castro, “Advances in the treatment of Chagas disease: Promising new drugs, plants and targets,” Biomed. Pharmacother., vol. 142, p. 112020, Oct. 2021, doi: 10.1016/j.biopha.2021.112020.

[5] S. Jayawardhana et al., “Benznidazole treatment leads to DNA damage in Trypanosoma cruzi and the persistence of rare widely dispersed non-replicative amastigotes in mice,” PLOS Pathog., vol. 19, no. 11, p. e1011627, Nov. 2023, doi: 10.1371/journal.ppat.1011627.

[6] A. M. Mejia et al., “Benznidazole-Resistance in Trypanosoma cruzi Is a Readily Acquired Trait That Can Arise Independently in a Single Population,” J. Infect. Dis., vol. 206, no. 2, pp. 220–228, Jul. 2012, doi: 10.1093/infdis/jis331.

[7] B. S. Hall, C. Bot, and S. R. Wilkinson, “Nifurtimox Activation by Trypanosomal Type I Nitroreductases Generates Cytotoxic Nitrile Metabolites,” J. Biol. Chem., vol. 286, no. 15, pp. 13088–13095, Apr. 2011, doi: 10.1074/jbc.M111.230847.

[8] G. I. Lepesheva, F. Villalta, and M. R. Waterman, “Targeting Trypanosoma cruzi Sterol 14α-Demethylase (CYP51),” in Advances in Parasitology, vol. 75, Elsevier, 2011, pp. 65–87. doi: 10.1016/B978-0-12-385863-4.00004-6.

[9] L. Friggeri et al., “Structural Basis for Rational Design of Inhibitors Targeting *Trypanosoma cruzi* Sterol 14α-Demethylase: Two Regions of the Enzyme Molecule Potentiate Its Inhibition,” J. Med. Chem., vol. 57, no. 15, pp. 6704–6717, Aug. 2014, doi: 10.1021/jm500739f.

[10] L. M. MacLean, J. Thomas, M. D. Lewis, I. Cotillo, D. W. Gray, and M. De Rycker, “Development of Trypanosoma cruzi in vitro assays to identify compounds suitable for progression in Chagas’ disease drug discovery,” PLoS Negl. Trop. Dis., vol. 12, no. 7, p. e0006612, Jul. 2018, doi: 10.1371/journal.pntd.0006612.

[11] I. Molina et al., “Randomized Trial of Posaconazole and Benznidazole for Chronic Chagas’ Disease,” N. Engl. J. Med., vol. 370, no. 20, pp. 1899–1908, May 2014, doi: 10.1056/NEJMoa1313122.

[12] D. Ortiz et al., “Targeting the Cytochrome *bc*_1_ Complex of Leishmania Parasites for Discovery of Novel Drugs,” Antimicrob. Agents Chemother., vol. 60, no. 8, pp. 4972–4982, Aug. 2016, doi: 10.1128/AAC.00850-16.

[13] S. Khare et al., “Utilizing Chemical Genomics to Identify Cytochrome b as a Novel Drug Target for Chagas Disease,” PLOS Pathog., vol. 11, no. 7, p. e1005058, Jul. 2015, doi: 10.1371/journal.ppat.1005058.

[14] A. H. Fairlamb and S. Wyllie, “The critical role of mode of action studies in kinetoplastid drug discovery,” Front. Drug Discov., vol. 3, p. 1185679, May 2023, doi: 10.3389/fddsv.2023.1185679.

[15] R. J. Wall et al., “The Q_i_ Site of Cytochrome *b* is a Promiscuous Drug Target in *Trypanosoma cruzi* and *Leishmania donovani*,” ACS Infect. Dis., vol. 6, no. 3, pp. 515–528, Mar. 2020, doi: 10.1021/acsinfecdis.9b00426.

[16] S. Khare et al., “Proteasome inhibition for treatment of leishmaniasis, Chagas disease and sleeping sickness,” Nature, vol. 537, no. 7619, pp. 229–233, Sep. 2016, doi: 10.1038/nature19339.

[17] A. Nagle et al., “Discovery and Characterization of Clinical Candidate LXE408 as a Kinetoplastid-Selective Proteasome Inhibitor for the Treatment of Leishmaniases,” J. Med. Chem., vol. 63, no. 19, pp. 10773–10781, Oct. 2020, doi: 10.1021/acs.jmedchem.0c00499.

[18] P. A. M. Michels, F. Bringaud, M. Herman, and V. Hannaert, “Metabolic functions of glycosomes in trypanosomatids,” Biochim. Biophys. Acta BBA - Mol. Cell Res., vol. 1763, no. 12, pp. 1463–1477, Dec. 2006, doi: 10.1016/j.bbamcr.2006.08.019.

[19] R. L. Berens, J. J. Marr, S. W. LaFon, and D. J. Nelson, “Purine metabolism in Trypanosoma cruzi,” Mol. Biochem. Parasitol., vol. 3, no. 3, pp. 187–196, Jul. 1981, doi: 10.1016/0166-6851(81)90049-9.

[20] J. D. N. Faria et al., “Inhibition of L-threonine dehydrogenase from Trypanosoma cruzi reduces glycine and acetate production and interferes with parasite growth and viability,” J. Biol. Chem., vol. 301, no. 2, p. 108080, Feb. 2025, doi: 10.1016/j.jbc.2024.108080.

[21] H. Acosta et al., “Proteomic analysis of glycosomes from Trypanosoma cruzi epimastigotes,” Mol. Biochem. Parasitol., vol. 229, pp. 62–74, Apr. 2019, doi: 10.1016/j.molbiopara.2019.02.008.

[22] Q. Li et al., “GMP synthase is essential for viability and infectivity of Trypanosoma brucei despite a redundant purine salvage pathway,” Mol. Microbiol., vol. 97, no. 5, pp. 1006–1020, Sep. 2015, doi: 10.1111/mmi.13083.

[23] L. Hedstrom, “IMP Dehydrogenase: Structure, Mechanism, and Inhibition,” Chem. Rev., vol. 109, no. 7, pp. 2903–2928, Jul. 2009, doi: 10.1021/cr900021w.

[24] D. R. Gollapalli, I. S. MacPherson, G. Liechti, S. K. Gorla, J. B. Goldberg, and L. Hedstrom, “Structural Determinants of Inhibitor Selectivity in Prokaryotic IMP Dehydrogenases,” Chem. Biol., vol. 17, no. 10, pp. 1084–1091, Oct. 2010, doi: 10.1016/j.chembiol.2010.07.014.

[25] L. Hedstrom, G. Liechti, J. B. Goldberg, and D. R. Gollapalli, “The antibiotic potential of prokaryotic IMP dehydrogenase inhibitors,” Curr. Med. Chem., vol. 18, no. 13, pp. 1909–1918, 2011, doi: 10.2174/092986711795590129.

[26] K. Mandapati et al., “Repurposing Cryptosporidium Inosine 5′-Monophosphate Dehydrogenase Inhibitors as Potential Antibacterial Agents,” ACS Med. Chem. Lett., vol. 5, no. 8, pp. 846–850, Aug. 2014, doi: 10.1021/ml500203p.

[27] N. N. Umejiego et al., “Targeting a Prokaryotic Protein in a Eukaryotic Pathogen: Identification of Lead Compounds against Cryptosporidiosis,” Chem. Biol., vol. 15, no. 1, pp. 70–77, Jan. 2008, doi: 10.1016/j.chembiol.2007.12.010.

[28] B. Striepen et al., “Genetic complementation in apicomplexan parasites,” Proc. Natl. Acad. Sci., vol. 99, no. 9, pp. 6304–6309, Apr. 2002, doi: 10.1073/pnas.092525699.

[29] J. E. Silverman Kitchin, M. K. Pomeranz, G. Pak, K. Washenik, and J. L. Shupack, “Rediscovering mycophenolic acid: A review of its mechanism, side effects, and potential uses,” J. Am. Acad. Dermatol., vol. 37, no. 3, pp. 445–449, Sep. 1997, doi: 10.1016/S0190-9622(97)70147-6.

[30] E. M. Eugui, A. Mirkovich, and A. C. Allison, “Lymphocyte-Selective Antiproliferative and Immunosuppressive Effects of Mycophenolic Acid in Mice,” Scand. J. Immunol., vol. 33, no. 2, pp. 175–183, Feb. 1991, doi: 10.1111/j.1365-3083.1991.tb03747.x.

[31] L. M. Shaw, M. Korecka, R. Venkataramanan, L. Goldberg, R. Bloom, and K. L. Brayman, “Mycophenolic Acid Pharmacodynamics and Pharmacokinetics Provide a Basis for Rational Monitoring Strategies,” Am. J. Transplant., vol. 3, no. 5, pp. 534–542, May 2003, doi: 10.1034/j.1600-6143.2003.00079.x.

[32] H. Park, “The emergence of mycophenolate mofetilin dermatology: from its roots in the world of organ transplantation to its versatile role in the dermatology treatment room,” J. Clin. Aesthetic Dermatol., vol. 4, no. 1, pp. 18–27, Jan. 2011.

[33] F. Bacal et al., “Mychophenolate Mofetil Increased Chagas Disease Reactivation in Heart Transplanted Patients: Comparison Between Two Different Protocols,” Am. J. Transplant., vol. 5, no. 8, pp. 2017–2021, Aug. 2005, doi: 10.1111/j.1600-6143.2005.00975.x.

[34] H. S. Oz, W. T. Hughes, E. K. Thomas, and C. J. McClain, “Effects of immunomodulators on acute Trypanosoma Cruzi infection in mice,” Med. Sci. Monit. Int. Med. J. Exp. Clin. Res., vol. 8, no. 6, pp. BR208-211, Jun. 2002.

[35] H. S. Te, G. Randall, and D. M. Jensen, “Mechanism of action of ribavirin in the treatment of chronic hepatitis C,” Gastroenterol. Hepatol., vol. 3, no. 3, pp. 218–225, Mar. 2007.

[36] J. Jain et al., “Characterization of Pharmacological Efficacy of VX-148, a New, Potent Immunosuppressive Inosine 5′-Monophosphate Dehydrogenase Inhibitor,” J. Pharmacol. Exp. Ther., vol. 302, no. 3, pp. 1272–1277, Jan. 2002, doi: 10.1124/jpet.102.035659.

[37] M. Huang, Y. Ji, K. Itahana, Y. Zhang, and B. Mitchell, “Guanine nucleotide depletion inhibits pre-ribosomal RNA synthesis and causes nucleolar disruption,” Leuk. Res., vol. 32, no. 1, pp. 131–141, Jan. 2008, doi: 10.1016/j.leukres.2007.03.025.

[38] D. Floryk and T. C. Thompson, “Antiproliferative effects of AVN944, a novel inosine 5-monophosphate dehydrogenase inhibitor, in prostate cancer cells,” Int. J. Cancer, vol. 123, no. 10, pp. 2294–2302, Nov. 2008, doi: 10.1002/ijc.23788.

[39] J. Jain, S. J. Almquist, D. Shlyakhter, and M. W. Harding, “VX-497: A novel, selective IMPDH inhibitor and immunosuppressive agent,” J. Pharm. Sci., vol. 90, no. 5, pp. 625–637, May 2001, doi: 10.1002/1520-6017(200105)90:5<625::AID-JPS1019>3.0.CO;2-1.

[40] X. Tong et al., “Merimepodib, an IMPDH inhibitor, suppresses replication of Zika virus and other emerging viral pathogens,” Antiviral Res., vol. 149, pp. 34–40, Jan. 2018, doi: 10.1016/j.antiviral.2017.11.004.

[41] W. Markland, T. J. McQuaid, J. Jain, and A. D. Kwong, “Broad-Spectrum Antiviral Activity of the IMP Dehydrogenase Inhibitor VX-497: a Comparison with Ribavirin and Demonstration of Antiviral Additivity with Alpha Interferon,” Antimicrob. Agents Chemother., vol. 44, no. 4, pp. 859–866, Apr. 2000, doi: 10.1128/AAC.44.4.859-866.2000.

[42] K. Ichinose et al., “Efficacy and Safety of Mizoribine by One Single Dose Administration for Patients with Rheumatoid Arthritis,” Intern. Med., vol. 49, no. 20, pp. 2211–2218, 2010, doi: 10.2169/internalmedicine.49.3810.

[43] I. Peña et al., “New Compound Sets Identified from High Throughput Phenotypic Screening Against Three Kinetoplastid Parasites: An Open Resource,” Sci. Rep., vol. 5, no. 1, p. 8771, Mar. 2015, doi: 10.1038/srep08771.

[44] D. Bajusz, A. Rácz, and K. Héberger, “Why is Tanimoto index an appropriate choice for fingerprint-based similarity calculations?,” J. Cheminformatics, vol. 7, no. 1, p. 20, Dec. 2015, doi: 10.1186/s13321-015-0069-3.

[45] R. Bentley, “Mycophenolic Acid: A One Hundred Year Odyssey from Antibiotic to Immunosuppressant,” Chem. Rev., vol. 100, no. 10, pp. 3801–3826, Oct. 2000, doi: 10.1021/cr990097b.

[46] Greg Landrum et al., rdkit/rdkit: 2025_03_1 (Q1 2025) Release. (Mar. 31, 2025). Zenodo. doi: 10.5281/ZENODO.591637.

[47] K. K. L. B. Adikaram, M. A. Hussein, M. Effenberger, and T. Becker, “Data Transformation Technique to Improve the Outlier Detection Power of Grubbs’ Test for Data Expected to Follow Linear Relation,” J. Appl. Math., vol. 2015, pp. 1–9, 2015, doi: 10.1155/2015/708948.

[48] S. Altschul, “Gapped BLAST and PSI-BLAST: a new generation of protein database search programs,” Nucleic Acids Res., vol. 25, no. 17, pp. 3389–3402, Sep. 1997, doi: 10.1093/nar/25.17.3389.

[49] K. Nass et al., “In cellulo crystallization of Trypanosoma brucei IMP dehydrogenase enables the identification of genuine co-factors,” Nat. Commun., vol. 11, no. 1, p. 620, Jan. 2020, doi: 10.1038/s41467-020-14484-w.

[50] F. Madeira et al., “The EMBL-EBI Job Dispatcher sequence analysis tools framework in 2024,” Nucleic Acids Res., vol. 52, no. W1, pp. W521–W525, Jul. 2024, doi: 10.1093/nar/gkae241.

[51] X. Robert and P. Gouet, “Deciphering key features in protein structures with the new ENDscript server,” Nucleic Acids Res., vol. 42, no. W1, pp. W320–W324, Jul. 2014, doi: 10.1093/nar/gku316.

[52] G. F. Mercaldi, A. T. Ranzani, and A. T. Cordeiro, “Discovery of New Uncompetitive Inhibitors of Glucose-6-Phosphate Dehydrogenase,” SLAS Discov., vol. 19, no. 10, pp. 1362–1371, Dec. 2014, doi: 10.1177/1087057114546896.

[53] G. F. Mercaldi et al., “*Trypanosoma cruzi* Malic Enzyme Is the Target for Sulfonamide Hits from the GSK Chagas Box,” ACS Infect. Dis., vol. 7, no. 8, pp. 2455–2471, Aug. 2021, doi: 10.1021/acsinfecdis.1c00231.

[54] F. W. Studier, “Protein production by auto-induction in high-density shaking cultures,” Protein Expr. Purif., vol. 41, no. 1, pp. 207–234, May 2005, doi: 10.1016/j.pep.2005.01.016.

[55] U. K. Laemmli, “Cleavage of Structural Proteins during the Assembly of the Head of Bacteriophage T4,” Nature, vol. 227, no. 5259, pp. 680–685, Aug. 1970, doi: 10.1038/227680a0.

[56] B. Striepen et al., “Gene transfer in the evolution of parasite nucleotide biosynthesis,” Proc. Natl. Acad. Sci., vol. 101, no. 9, pp. 3154–3159, Mar. 2004, doi: 10.1073/pnas.0304686101.

[57] J. P. Richard, X. Zhai, and M. M. Malabanan, “Reflections on the catalytic power of a TIM-barrel,” Bioorganic Chem., vol. 57, pp. 206–212, Dec. 2014, doi: 10.1016/j.bioorg.2014.07.001.

[58] G. L. Prosise and H. Luecke, “Crystal Structures of Tritrichomonasfoetus Inosine Monophosphate Dehydrogenase in Complex with Substrate, Cofactor and Analogs: A Structural Basis for the Random-in Ordered-out Kinetic Mechanism,” J. Mol. Biol., vol. 326, no. 2, pp. 517–527, Feb. 2003, doi: 10.1016/S0022-2836(02)01383-9.

[59] X. Wang, P. Kuzmic, and L. Hedstrom, “Mechanism of Inhibitor Selectivity Revealed by Mutagenesis and pre- Steady-State Studies,” FASEB J., vol. 35, no. S1, p. fasebj.2021.35.S1.02852, May 2021, doi: 10.1096/fasebj.2021.35.S1.02852.

[60] T. Bessho et al., “Characterization of the novel *Trypanosoma brucei* inosine 5′-monophosphate dehydrogenase,” Parasitology, vol. 140, no. 6, pp. 735–745, May 2013, doi: 10.1017/S0031182012002090.

[61] F. Dobie, A. Berg, J. M. Boitz, and A. Jardim, “Kinetic characterization of inosine monophosphate dehydrogenase of Leishmania donovani,” Mol. Biochem. Parasitol., vol. 152, no. 1, pp. 11–21, Mar. 2007, doi: 10.1016/j.molbiopara.2006.11.007.

[62] P. W. Hager, F. R. Collart, E. Huberman, and B. S. Mitchell, “Recombinant human inosine monophosphate dehydrogenase type I and type II proteins,” Biochem. Pharmacol., vol. 49, no. 9, pp. 1323–1329, May 1995, doi: 10.1016/0006-2952(95)00026-V.

[63] T. Hishiki et al., “Identification of IMP Dehydrogenase as a Potential Target for Anti-Mpox Virus Agents,” Microbiol. Spectr., vol. 11, no. 4, pp. e00566–23, Aug. 2023, doi: 10.1128/spectrum.00566-23.

[64] G. Mei-jiao et al., “Antiviral effects of selected IMPDH and DHODH inhibitors against foot and mouth disease virus,” Biomed. Pharmacother., vol. 118, p. 109305, Oct. 2019, doi: 10.1016/j.biopha.2019.109305.

[65] D. Floryk and T. C. Thompson, “Antiproliferative effects of AVN944, a novel inosine 5-monophosphate dehydrogenase inhibitor, in prostate cancer cells,” Int. J. Cancer, vol. 123, no. 10, pp. 2294–2302, Nov. 2008, doi: 10.1002/ijc.23788.

[66] A. G. Zimmermann, J.-J. Gu, J. Laliberté, and B. S. Mitchell, “Inosine-5′-Monophosphate Dehydrogenase: Regulation of Expression and Role in Cellular Proliferation and T Lymphocyte Activation,” in Progress in Nucleic Acid Research and Molecular Biology, vol. 61, Elsevier, 1998, pp. 181–209. doi: 10.1016/S0079-6603(08)60827-2.

[67] G. Gao et al., “Single-cell RNA sequencing in double-hit lymphoma: IMPDH2 induces the progression of lymphoma by activating the PI3K/AKT/mTOR signaling pathway,” Int. Immunopharmacol., vol. 125, p. 111125, Dec. 2023, doi: 10.1016/j.intimp.2023.111125.

[68] R. B. Klisovic et al., “A phase I trial of AVN944 in patients with advanced hematologic malignancies,” J. Clin. Oncol., vol. 25, no. 18_suppl, pp. 14026–14026, Jun. 2007, doi: 10.1200/jco.2007.25.18_suppl.14026.

[69] S. Li, M. Gong, J. Shao, Y. Sun, Y. Zhang, and H. Chang, “Antiviral activity of merimepodib against foot and mouth disease virus in vitro and in vivo,” Mol. Immunol., vol. 114, pp. 226–232, Oct. 2019, doi: 10.1016/j.molimm.2019.07.021.

[70] J. J. Gu et al., “Inhibition of T lymphocyte activation in mice heterozygous for loss of the IMPDH II gene,” J. Clin. Invest., vol. 106, no. 4, pp. 599–606, Aug. 2000, doi: 10.1172/JCI8669.

[71] J. S. Dayton, T. Lindsten, C. B. Thompson, and B. S. Mitchell, “Effects of human T lymphocyte activation on inosine monophosphate dehydrogenase expression.,” J. Immunol., vol. 152, no. 3, pp. 984–991, Feb. 1994, doi: 10.4049/jimmunol.152.3.984.

[72] S. J. Calise, G. Abboud, H. Kasahara, L. Morel, and E. K. L. Chan, “Immune Response-Dependent Assembly of IMP Dehydrogenase Filaments,” Front. Immunol., vol. 9, p. 2789, Nov. 2018, doi: 10.3389/fimmu.2018.02789.

[73] J. De Meis, A. Morrot, D. A. Farias-de-Oliveira, D. M. S. Villa-Verde, and W. Savino, “Differential Regional Immune Response in Chagas Disease,” PLoS Negl. Trop. Dis., vol. 3, no. 7, p. e417, Jul. 2009, doi: 10.1371/journal.pntd.0000417.

[74] A. Morrot, J. Barreto De Albuquerque, L. R. Berbert, C. E. De Carvalho Pinto, J. De Meis, and W. Savino, “Dynamics of Lymphocyte Populations during *Trypanosoma cruzi* Infection: From Thymocyte Depletion to Differential Cell Expansion/Contraction in Peripheral Lymphoid Organs,” J. Trop. Med., vol. 2012, pp. 1–7, 2012, doi: 10.1155/2012/747185.

[75] F. F. De Araújo et al., “Regulatory T Cells Phenotype in Different Clinical Forms of Chagas’ Disease,” PLoS Negl. Trop. Dis., vol. 5, no. 5, p. e992, May 2011, doi: 10.1371/journal.pntd.0000992.

[76] M. A. Bryan, S. E. Guyach, and K. A. Norris, “Specific Humoral Immunity versus Polyclonal B Cell Activation in Trypanosoma cruzi Infection of Susceptible and Resistant Mice,” PLoS Negl. Trop. Dis., vol. 4, no. 7, p. e733, Jul. 2010, doi: 10.1371/journal.pntd.0000733.

[77] L. S. A. Passos et al., “Activation of Human CD11b+ B1 B-Cells by Trypanosoma cruzi-Derived Proteins Is Associated With Protective Immune Response in Human Chagas Disease,” Front. Immunol., vol. 9, p. 3015, Jan. 2019, doi: 10.3389/fimmu.2018.03015.

[78] E. Pérez-Antón et al., “Impact of benznidazole treatment on the functional response of Trypanosoma cruzi antigen-specific CD4+CD8+ T cells in chronic Chagas disease patients,” PLoS Negl. Trop. Dis., vol. 12, no. 5, p. e0006480, May 2018, doi: 10.1371/journal.pntd.0006480.

[79] R. Naffouje et al., “Anti-Tumor Potential of IMP Dehydrogenase Inhibitors: A Century-Long Story,” Cancers, vol. 11, no. 9, p. 1346, Sep. 2019, doi: 10.3390/cancers11091346.

